# Data-driven Topological Filtering based on Orthogonal Minimal Spanning Trees: Application to Multi-Group MEG Resting-State Connectivity

**DOI:** 10.1101/187252

**Authors:** Stavros I. Dimitriadis, Marios Antonakakis, Panagiotis Simos, Jack M. Fletcher, Andrew C. Papanicolau

## Abstract

In the present study a novel data-driven topological filtering technique is introduced to derive the backbone of functional brain networks relying on orthogonal minimal spanning trees (OMST). The method aims to identify the essential functional connections to ensure optimal information flow via the objective criterion of global efficiency minus the cost of surviving connections. The OMST technique was applied to multichannel, resting-state neuromagnetic recordings from four groups of participants: healthy adults (n=50), adults who have suffered mild traumatic brain injury (n=30), typically developing children (n=27), and reading-disabled children (n=25). Weighted interactions between network nodes (sensors) were computed using an integrated approach of dominant intrinsic coupling modes based on two alternative metrics (symbolic mutual information and phase lag index), resulting in excellent discrimination of individual cases according to their group membership. Classification results using OMST-derived functional networks were clearly superior to results using either relative power spectrum features or functional networks derived through the conventional minimal spanning trees algorithm.

## Introduction

Neuronal populations generate oscillatory electrical activity as a result of complex neurophysiological processes taking place within individual neurons and across neuronal populations (Buzsaki et al., 2004, 2006; Llinas, 2014). Such firing patterns can give rise to synchronized input to other cortical areas, supporting the interaction of a given assembly with more distant neuronal assemblies at the prominent oscillating frequency of the source population (Shew et al., 2009). It has been proposed that this cross-frequency coupling (CFC) promotes accurate timing between different oscillatory rhythms and dynamic control of distributed functional networks (Canolty and Knight, 2010; Varela et al., 2001; Buzsaki, 2006). Magnetoencephalography (MEG) is uniquely suited to address functional connectivity based on CFC because it possesses adequate temporal resolution to describe the real-time dynamics of fine-grained interactions between neuronal populations. There is rapidly accumulating experimental evidence supporting the role of CFC in cognition (Canolty and Knight, 2010; Buzsaki and Watson 2012; Jirsa and Muller, 2013; Dimitriadis et al., 2015a,c, 2016) and as a marker of neurophysiological dysfunction in developmental disorders such as reading disability (Dimitriadis et al., 2016b).

Neuronal interactions at the basic brain rhythms (Buzsaki, 2006) can be quantified through a variety of connectivity estimators, each featuring distinct advantages and limitations (Bastos and Schoffelen, 2016). The application of any type of connectivity estimator to a multichannel recording set leads to a fully-connected graph containing a large proportion of potentially spurious connections. Identifying such spurious interactions requires statistical filtering (Aru et al., 2014). The most common approach toward this goal is through surrogate analysis that permits calculation of p-values associated with each interaction, which are then thresholded using an adaptive criterion, such as False Discovery Rate, to control for Type I error.

Following statistical filtering, surviving interactions typically need to undergo spatial (topological) filtering in order to derive a network structure that contains only the essential interactions between nodes and is consequently more likely to be meaningful from a neuroscience perspective (Bullmore and Bassett, 2011; Van Wijk et al., 2010). Existing topological filtering approaches rely on, largely, arbitrary criteria, such as absolute weight threshold (e.g., > 0.5), upper density limits (e.g., keeping the strongest 10% of connections), and mean graph degree (e.g., retaining connections so that the mean degree value is kept > 5; Dimitriadis et al., 2010). A recent study explored the caveats of applying proportional thresholding on fMRI resting-state brain networks from clinical populations (van den Heuvel et al., 2017). The aforementioned observations highlight the need for *data-driven* topological filtering techniques. In principle, the latter may possess greater sensitivity to network features to serve as connectomic biomarkers for disorders such as Alzheimer’s disease, schizophrenia, autism, and reading disability. Data-driven techniques are also crucial to ensure compatibility of results across laboratories and/or scanner types where absolute threshold criteria are not applicable (Abraham et al., 2017; Dansereau et al., 2017).

An increasingly popular, assumption-free method for identifying the essential set of connections within a fully connected graph is based on Minimal Spanning Trees (MST; Meier et al., 2015; Tewarie et al., 2015). More specifically, the MST connects all the N nodes in a graph through N-1 connections my minimizing the total cost of information flow and without introducing cycles. The method addresses crucial limitations of existing topological filtering schemes, which rely on absolute threshold or density and, additionally, preserves the connectedness of the brain network. However, the conventional MST approach typically results in trees with only N-1 links, which for large graphs are too sparse to allow reliable discrimination between two (Dimitriadis et al., 2015a; Antonakakis et al., 2016) or more groups (Khazaeea et al., 2016). To address this problem, the orthogonal MST approach (OMST) was introduced (Dimitriadis et al., 2017) by utilizing alternative algorithms to construct the MST of a weighted graph (Kruskal, 1956; Prim, 1957). The OMST method preserves the main advantage of MST (i.e., assumption-free, data-driven approach that maintains network connectedness) and further ensures a denser and, potentially, more meaningful network. It is implemented by sampling connections over multiple rounds of MST in order to identify the subset of functional interactions that would ensure optimal information flow (indexed by network global efficiency) while minimizing the cost incurred by preserved functional connections. OMST has been used in pattern recognition and computer vision task as a re-ranking method (Fotopoulou et al., 2014). The superior performance of this topological filtering approach over several conventional filtering schemes has recently been demonstrated using large EEG and fMRI databases (Dimitriadis et al., 2017).

In the present work we demonstrate the advantages of OMST as a topological filtering approach for sensor-level, resting-state neuromagnetic recordings. At the temporal scale characteristic of CFC, source localization (and related arbitrary choices of algorithms and anatomic templates) may introduce significant distortions to the source-level (reconstructed) signals. This added layer of complexity, although in principle desirable to enhance the anatomic relevance of results, would likely have confounded the primary goal of the study.

In addition to using OMST, a novel feature of the present work involves use of mutual information derived from symbolized time series (symbolic mutual information; SMI) to quantify the strength of coupling between MEG sensors both within-and between-predefined frequency bands (i.e., cross-frequency coupling; Robinson and Mandell, 2015). In this approach, neuromagnetic signals are first transformed into symbolic sequences consisting of a finite set of substrings (Janson et al., 2004). Signal complexity was assessed by the degree of repeatability of substring sequences over time using the symbolization procedure described in Dimitriadis et al. (2016a). The theoretical advantage of SMI lies in its capacity to represent each pair of time series as a set of two symbolic sequences utilizing a *common* set of symbols. SMI is a weighted connectivity estimator which describes interactions between any two signals in the form of the strength of linear and non-linear functional associations (King et al., 2013; Robinson and Mandell, 2015). Being less susceptible to artifacts, SMI was chosen to handle MEG data from young children in the current study. Moreover, SMI was favored over delay Transfer Entropy, which may be more appropriate for source-level data (Roux et al., 2013). A more conventional connectivity estimator (Phase Lag Index; PLI) was also employed to derive functional connectivity graphs which were submitted to the OMST-based topological filtering in a separate analysis.

Briefly the analysis pipeline adopted in the present study involves the use of surrogate data sets to perform statistical filtering of functional connections, resulting in integrated functional brain networks featuring the dominant types of sensor interactions for each participant (Engel et al., 2013; Dimitriadis et al., 2016b). Such sparse networks were obtained independently for SMI and PLI and were subsequently filtered, topologically, using OMST. The sensitivity of this procedure to differences in resting-state brain connectivity attributed to participant age, presence of reading disability, and history of acute brain insult (mild traumatic brain injury; MTBI) was assessed on a large dataset consisting of four subgroups of participants: healthy adults (n=50), adults who have suffered MTBI (n=30), typically developing children (n=27), and reading-disabled children (n=25).

## Materials and Methods

### Participants

For the demonstration of the proposed algorithm, we used resting-state neuromagnetic recordings from four groups: healthy adults (n=50; 31 women, aged: 33.5±9.32 years with 15.4 ± 3.3 years of formal education), adults who had suffered mild traumatic brain injury (n=30; 13 women, aged: 32.3±9.9 years with 15.1 ± 2.9 years of formal education), typically developing children (n=27; 15 girls, aged: 10.45±2.6 years), and reading-disabled children (n=25; 14 girls, aged: 11.05±2.42 years). Resting state data were collected as part of ongoing projects at the Magnetoencephalography Laboratory, University of Texas Health Science Center-Houston. Detailed information on particpant characteristics can be found elsewhere (Antonakakis et al., 2016; Dimitriadis et al., 2013, 2015, 2016b).

### Preprocessing

The MEG data underwent artifact reduction using Matlab (The MathWorks, Inc., Natick, MA, USA) and Fieldtrip (Oostenveld et al., 2011). Independent component analysis (ICA) was used to separate cerebral from non-cerebral activity using the extended Infomax algorithm as implemented in EEGLAB (Delorme and Makeig, 2004). The data were whitened and reduced in dimensionality using principal component analysis with a threshold set to 95% of the total variance (Delorme and Makeig, 2004; Escudero et al., 2011). Kurtosis, Rényi entropy, and skewness values of each independent component were used to identify and remove ocular and cardiac artifacts. A given component was considered an artifact if, after normalization to zero mean and unit variance, more than 20% of its values were greater/lower than 2 SDs from the mean (Escudero et al., 2011; Dimitriadis et al., 2013, 2015a,b; Antonakakis et al., 2015, 2016). To further ensure that Independent Components meeting the aforementioned criterion were indeed artifactual, we examined their time course and morphology (characteristic for cardiac and myogenic artifacts). In addition, source localization was performed using linearly constrained minimum variance beamformers (van Veen et al., 1997) to ensure that source locations at the magnetic field peak of each artifact were outside the brain.

Subsequently, the reconstructed axial gradiometer recordings were transformed into planar gradiometer field approximations using the *sincos* method implemented in Fieldtrip (Oostenveld et al., 2011). The data were finally bandpass-filtered in the following frequency ranges using a 3^rd^-order Butterworth filter (in zero-phase mode): 0.5-4, 4-8, 8-10, 10-13, 13-15, 15-19, 20-29, and 30-45Hz corresponding to δ, θ, (α_1_, α_2_, β_1_, β_2_, β_3_, and γ bands.

### Integrated Functional Connectivity Graphs

The strength of intra-and inter-frequency coupling for each pair of sensors was indexed by the undirected, weighted SMI (King et al., 2013; Robinson and Mandell, 2015). Initially, each pair of time series was transformed into two symbolic sequences utilizing a *common* set of symbols using the Neural Gas algorithm, which was first adapted to handle time series pairs (see Section 1 in Supplementary Material). Our group has demonstrated the utility and relative advantages of the Neural Gas algorithm in identifying dynamic functional graphs and introduced the notion of functional connectivity microstates (Dimitriadis et al., 2013). Additionally, we have used the Neural Gas algorithm to symbolize pairs of time series and then estimate delay symbolic transfer entropy (Dimitriadis et al., 2016a). We have further demonstrated that the Neural Gas algorithm produced more stable results which were proven more robust to various types of noise, compared to ordinal pattern analysis, a frequently employed alternative method to symbolize time series.

SMI is defined as:

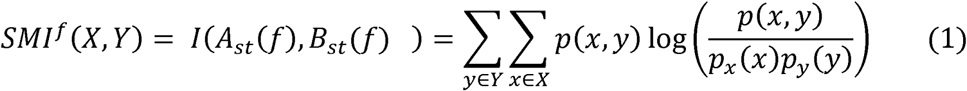

where X = ^A^s_t_ and Y = ^B^s_t_ are the two symbolic sequences, p(x,y) is the joint probability distribution function of X and Y, and p_x_(x) = ∑_y∈y_p(x,y) and p_y_(y) = ∑_x∈X_p(x,y) are the marginal probability distribution functions of X and Y, respectively. SMI values range between 0 and 1, with 0 denoting no functional coupling and 1 indicating perfect functional coupling over the entire recording period. This procedure resulted in a single functional connectivity graph (FCG) per participant, frequency band (8), and pair of frequency bands (28) consisting of SMI values.

Individual FCGs were submitted to statistical filtering using surrogate data to determine the Dominant Intrinsic Coupling Mode (DICM) for each pair of symbolic sequences (sensors). 10,000 surrogate data sets were created by shuffling the symbolic sequence of the second MEG sequence (^B^s_t_) in each pair (^A^s_t_, ^B^s_t_) and reestimated the SMI values. The concept of DICM is closely linked to the notion that although the specific frequencies and strengths of interactions between sensors may vary during the resting-state recording for a given participant, each sensor pair displays a typical (i.e., more temporally stable) mode of interaction which can identified via application of a conservative statistical criterion using surrogate data (Dimitriadis et al., 2016b,c). In the present work, a p-value was assigned to each pair of symbolic sequences (same-frequency/between-sensor, cross-frequency/between-sensor, and cross-frequency/within-sensor pairs) reflecting the proportion of permutations that yielded surrogate SMI values higher than the observed SMI values. This procedure produced a 3D tensor of p values for each participant of size 36 × 248 × 248. Significant DICM(s) for each pair of symbolic sequences were determined by applying a Bonferroni-adjusted p < 0.01/36 = 0.00028 in order to control for family-wise Type I error. When more than one frequency or frequency pairs exceeded this threshold, the one associated with the lowest p value was retained. This procedure resulted in two 2D matrices for each participant of size 248 × 248: one containing the highest/statistically significant SMI values and the second the identity of the corresponding frequency or frequency pair (e.g., 1 for δ, 2 for θ, …, 8 for γ, 9 for δ-θ, …, 15 for δ-γ,…, 36 for β_3_-β).

For comparison, FCGs were also constructed using a conventional connectivity metric, Phase Lag Index (PLI; Stam et al., 2007), which is considered to be less susceptive to volume conduction (see Section 2 in Supplementary Material).

### Topological filtering of Functional Connectivity Graphs using OMST

A crucial difference of the Orthogonal Minimal Spanning Trees (OMST) algorithm from the conventional MST method is that the latter tends to preserve the weakest connections under the constraint of minimizing overall cost of connecting all the nodes in the graph. To address this limitation functional connectivity graphs were first inverted to emphasize the strongest connections corresponding to higher SMI values.

The proposed OMST algorithm was applied to the statistically thresholded FCGs, independently for each participant, as follows (Dimitriadis et al., 2017):

a. The MSTs were extracted by iteratively applying Kruskal’s algorithm on the inverted weighted Functional Connectivity Graphs containing the Dominant Intrinsic Coupling Mode for each pair of sensors.
b. After extracting the 1^st^ MST, which connects all the N sensors through N-1 edges, the N-1 edges were substituted with ‘Inf’ in the original network in order to avoid capturing the same edges and also to maintain orthogonality with the next MST. Then a 2^nd^ MST was estimated that connects all of the N sensors with minimal total distance, satisfying the constraint that it is orthogonal (i.e., does not share common edges) with the 1^st^ MST. Next, the N-1 connections of the 2^nd^ MST were substituted with zeros and a 3^rd^ MST was estimated that connected the sensors with the minimal total weight, subject to the constraint that it is orthogonal to the previous two constructed MST’s (1^st^ and 2^nd^). In general, an m^th^-MST is orthogonal to all the previous (m-1)^th^ MSTs, having exactly m(N-1) edges.
c. Connections were aggregated across OMSTs (including the 1^st^) in order to optimize the function of global efficiency (GE) minus Cost over Cost as described in more detail in (e) below. For instance, this step can aggregate 3*(N-1) edges from the first three OMSTs plus the 4^th^ MST.
d. For each added connection to the aggregated network, the objective function of Global Cost Efficiency (GCE) = GE-Cost was estimated, where Cost denotes the ratio of the total weight of the selected edges, over multiple iterations of OMST, divided by the total strength of the original fully-weighted graph. The values of this formula range within the limits of an economical small-world network for healthy control participants (Bassett and Bullmore, 2006). The network which is considered as functionally optimal is the one associated with the maximum value of the following quality formula:

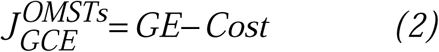
e. Topological filtering on the person-specific FCGs featuring the Dominant Intrinsic Coupling Modes entailed retaining those sensor interactions that optimized the function of global efficiency (GE) minus Cost over Cost. A sample plot of this function from a typical reader, obtained after running exhaustive OMSTs until all observed weights were tested, is shown in Fig. 1. The maximum of this (always) positive curve reflects the optimization of the proposed OMST algorithm. In the example of Fig. 1, the GE-Cost vs. Cost function was optimized after 11 OMSTs leading to a selection of 2,689 connections—a mere 8.9% of the total number of connections which were initially retained following statistical filtering.

**Fig. 1.**
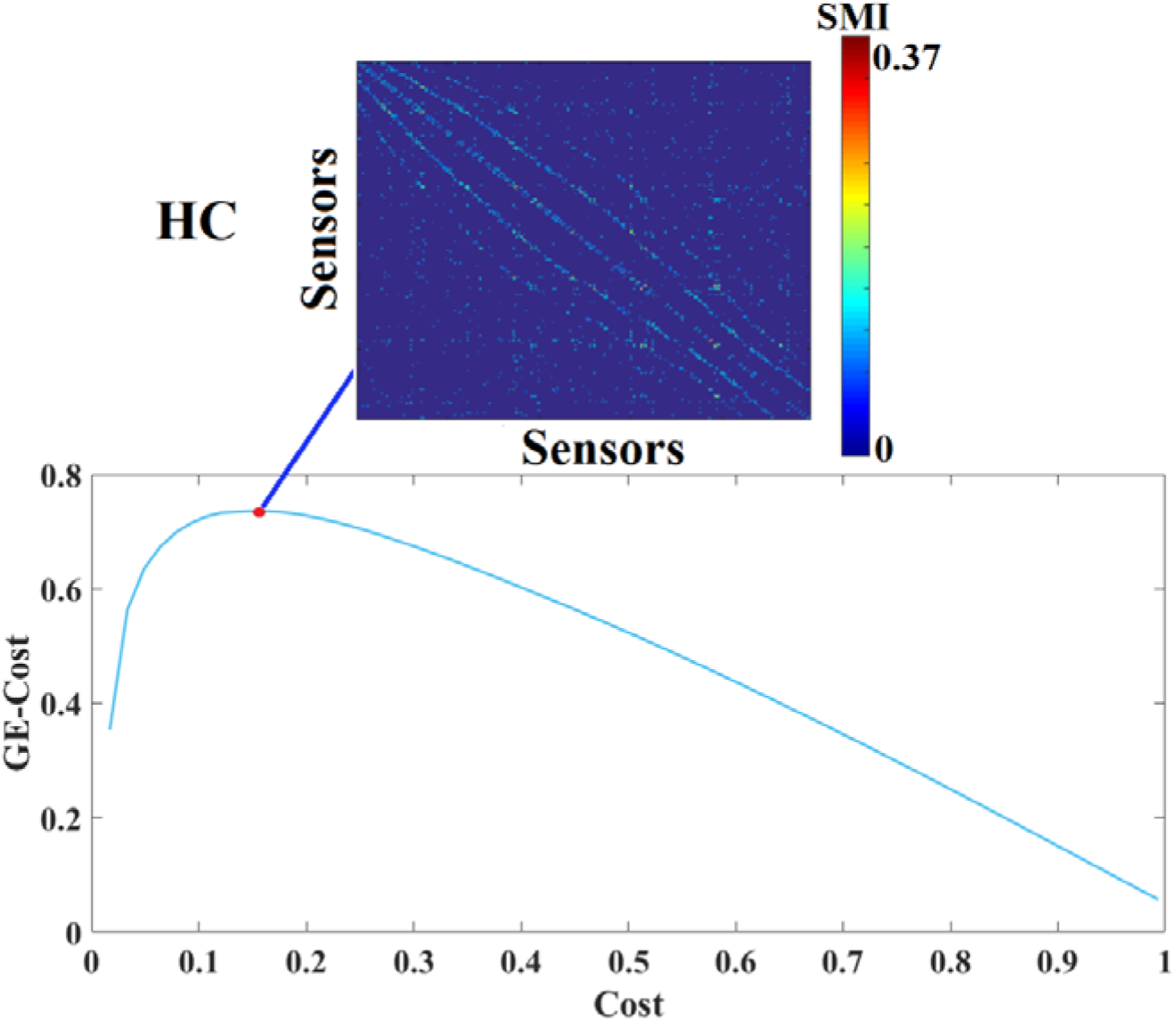
Optimization of the function of Global Efficiency (GE) minus Cost over Cost over multiple OMST runs in data from a typical reader. The red circle denotes the peak of the computed curve, while the resulting topologically filtered Functional Connectivity Graphs (FCG) is shown in the inset. Abbreviation; SMI: Symbolic Mutual Information. HC: Typically developing children,

### Person-specific graph metrics of topologically filtered Functional Connectivity Graphs

Subject-specific, OMST-filtered FCGs were characterized using the following network metrics: global efficiency (GE), eccentricity, radius, and diameter. Global efficiency is the average inverse shortest path length in the network and is, perhaps, is the most informative estimator of the integration of information flow within a network. Eccentricity is defined as the maximum shortest path length between a given sensor and any other sensor, whereas the radius and diameter correspond to the average and maximum eccentricity values across all sensors, respectively. Graph metrics selected for the present study represent the most widely used across imaging modalities and research questions (Bullmore and Sporns, 2009; Rubinov and Sporns, 2010; Telesford et al., 2011; Stam, 2015). Pairwise group comparisons on each network metric were performed using the Wilcoxon-Rank Sum Test (evaluated at a conservative p < 0.0001).

### Graph Diffussion Distance: A metric of group differences on network structure

In order to assess group differences in the OMST-filtered Functional Connectivity Graphs at the single-case level, we computed the Graph Diffusion Distance metric (Fouss et al., 2012; Hammond et al., 2013). The graph laplacian operator of each subject-specific FCG was defined as L = D – FCG, where D is a diagonal degree matrix related to FCG. This method entails modeling hypothetical patterns of information flow among sensors based on each observed (static) FCG. The diffusion process on the person-specific FCG was allowed for a set time t; the quantity that underwent diffusion at each time point is represented by the time-varying vector *u(t)* ∈ ℜ^*N*^. Thus, for a pair of sensors i and j, the quantity FCGij (ui(t) – uj(t)) represents the hypothetical flow of information from i to j via the edges that connect them (both directly and indirectly). Summing all these hypothetical interactions for each sensor leads to 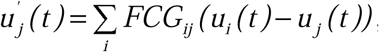 which can be written as:

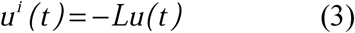

where L is the graph laplacian of FCG. At time t = 0 Equation 2 has the analytic solution: *u(t)=exp(-tL)u^(0)^*. Here exp(-tL) is a N × N matrix function of t, known as Laplacian exponential diffusion kernel (Fouss et al., 2012), and u^(0)^ = e_j_, where *e*_*j*_∈ℜ ^*N*^ is the unit vector with all zeros except in the j^th^ component. Running the diffusion process through time t produced the diffusion pattern exp(-tL)ej which corresponds to the jth column of exp(-tL).

Next, a metric of dissimilarity between every possible pair of person-specific diffusion-kernelized FCGs (FCG1,FCG2) was computed in the form of the graph diffusion distance d_gdd_(t). The higher the value of d_g_dd(t) between two graphs, the more distinct is their network topology as well as the corresponding, hypothetical information flow. The columns of the Laplacian exponential kernels, exp(-tL1) and exp(-tL2), describe distinct diffusion patterns, centered at two corresponding sensors within each FCG. The d_gdd_(t) function is searching for a diffusion time t that maximizes the Frobenius norm of the sum of squared differences between these patterns, summed over all sensors, and is computed as:

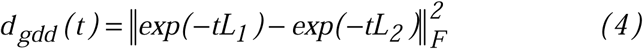

where 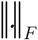 is the Frobenius norm.

Given the spectral decomposition L=VΛV (V defines the eigenvectors and Λ the eigenvalues), the laplacian exponential can be estimated via:

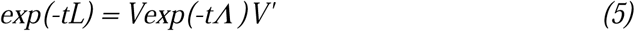

where for Λ, exp(-tΛ) is diagonal to the ith entry given by *e*^-*tΛ*_i,i_^. We computed d_gdd_(FCG_1_,FCG_2_) by first diagonalizing L1 and L2 and then applying equations (3) and (4) to estimate d_gdd_(t) for each time point t of the diffusion process. In this manner, a single dissimilarity value was computed for each pair of participants based on their corresponding FCGs.

### Group differences on relative spectral power

Relative power in each frequency band was examined as a lower-level feature that could account for group differences in FCGs. Initially, statistical filtering was to the RP values obtained for each of 8 frequency bands (Fr) and 248 sensors (S) by first computing Laplacian scores (LSFr_S). The null distribution for each of the 1984 features was obtained through bootstrapping by randomizing the group identity labels assigned to each feature for 10,000 times and estimating the corresponding Laplacian scores. Next, we assessed deviations of the Laplacian score of each feature from the null distribution 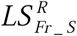 and assigned a (one-sided) p-value as the percentage of observed 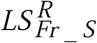 exceeding the original estimated LS_Fr_S_ (evaluated at a Bonferroni-corrected p < 0.05/(8*248)).

### Group separation and classification

A k-nearest neighbor classifier was employed to assess the accuracy of assigning cases to each of the four study groups based on either the Graph Diffusion Distance metric derived from the OMST-filtered functional connectivity graphs or the statistically filtered relative power metrics.

Results were obtained for two classification schemes to permit direct comparison with those obtained using the proposed OSMT-GDD scheme: a) contrasting groups in a pair-wise fashion and b) multi-group classification.

Finally, Graph Diffusion Distance values were projected to a common 3D space using multidimensional scaling, as a means of visualizing the level of similarity of individual cases (Borg and Groenen, 2005). The multidimensional scaling algorithm aims to place each case in *N*-dimensional space by preserving between-case distances. Each case is then assigned coordinates on each of a predetermined set of *N* dimensions (N=3 in the present work).

## Results

### Group characteristics on topologically filtered Functional Connectivity Graphs

Typically achieving students showed higher eccentricity and radius values, as well as smaller diameter values than adult typical readers (Fig. 2). Moreover, RD children showed lower global efficiency and higher eccentricity, radius, and diameter values than age-matched typical readers. Participants with a history of mTBI displayed higher diameter and lower eccentricity and higher global efficiency and diameter values than healthy adults.

**Fig. 2.**
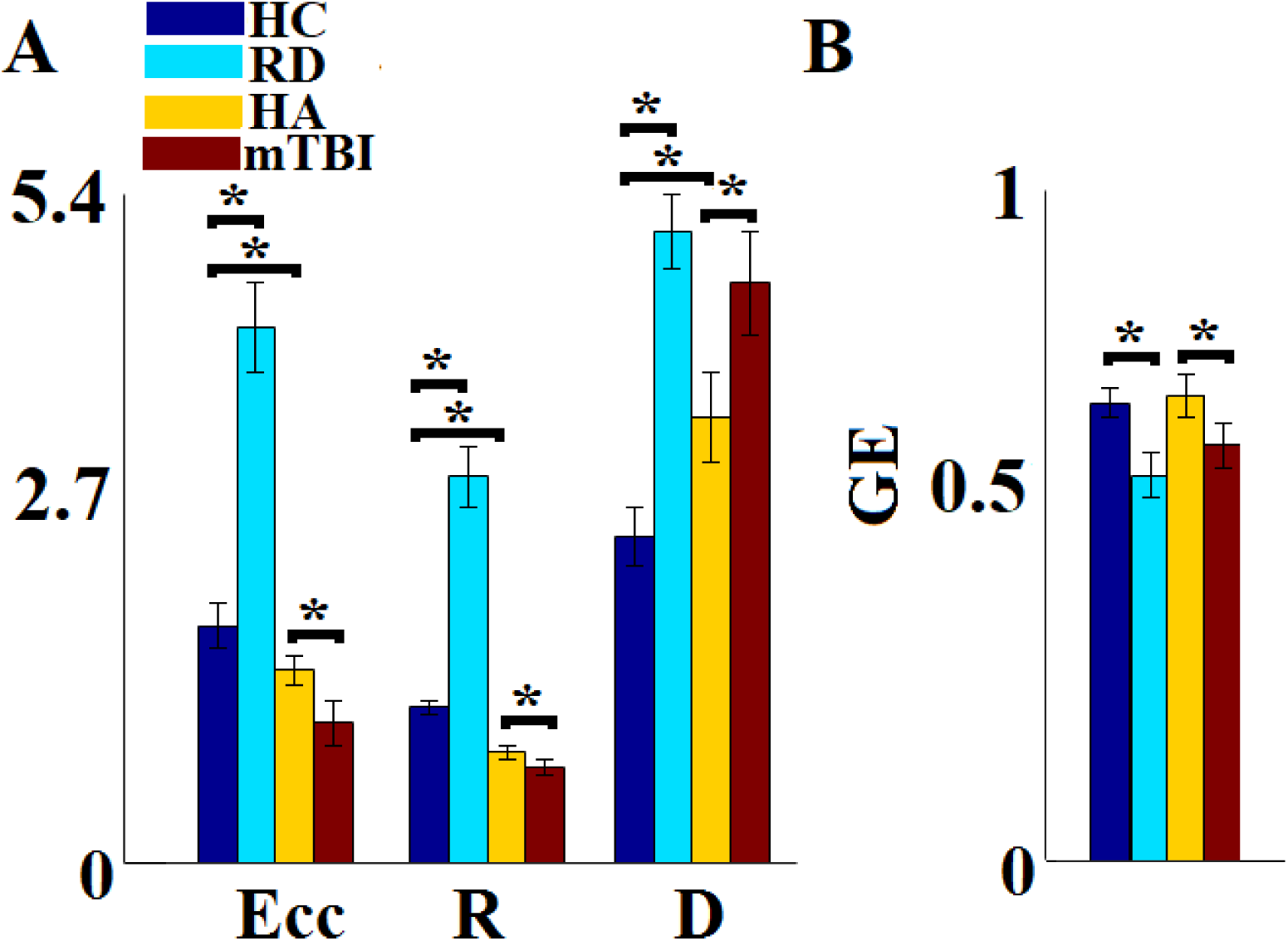
Group-averaged network metrics of the topologically filtered FCGs characterizing MEG resting-state data. Brackets indicate significant pair-wise group differences (Wilcoxon Rank Sum Test p < 0.0001). Abbreviations; HC: Typically developing children, RD: Reading-Disabled children, HA: Healthy Adults, mTBI: mild Traumatic Brain injury patients; GE: global efficiency, Ecc: Eccentricity, R: radius, D: diameter.

The reliability of OMST-derived network metrics was further assesed through split-half analyses, whereby global efficiency, eccentricity, radius, and diameter values were recomputed for each age-and gender-matched split half sections of the four study groups. As shown in Figure S2 (Supplementary Material) average network metrics were very similar between split-half subgroups (p> .15 in all cases).

### Group differences on OMST-derived network features

Applying d_gdd_(t) in a pair-wise fashion on person-specific FCGs based on Symbolic Mutual Information across the 132 participants produced the dissimilarity matrix displayed in Fig. 3A. The clear group separation was visualized by projecting individual Graph Diffusion Distance values onto a common 3D space following dimensionality reduction using multidimensional scaling (Fig. 3B). Classification accuracy reached 100% for both the pairwise and multi-group contrasts. As displayed in Fig. S1 (Supplementary Material) similar results were obtained for the OMST-derived PLI-based FCGs.

**Fig. 3.**
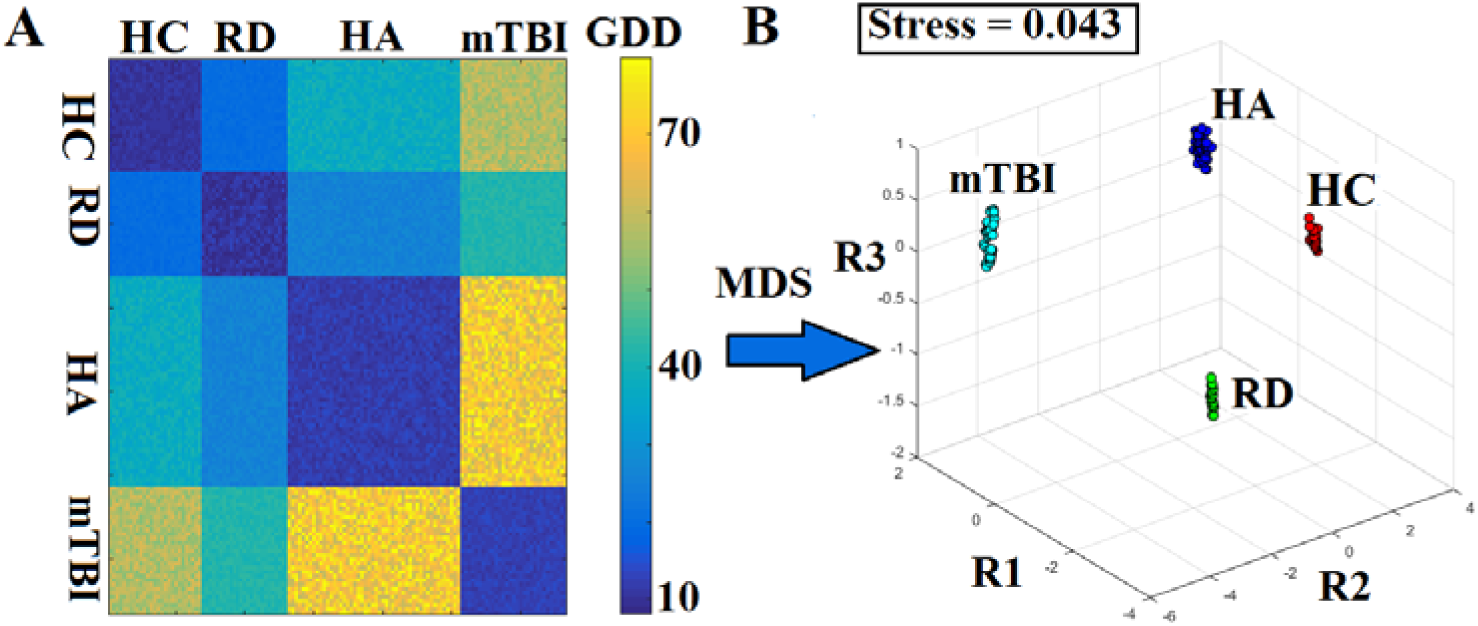
Topological filtering of Graph Diffusion Distance (GDD) values using Orthogonal MST. A) Dissimilarity matrix of subject-specific FCGs (N=132) based on the GDD metric. B) Multi-dimensional Scaling (MDS) was applied to the dissimilarity matrix of GDD values which were rescaled and projected in a common 3D space. Stress indicates the % loss of information due to the dimensionality reduction process via the MDS algorithm. Abbreviations; HC: Typically developing children, RD: Reading-Disabled children, HA: Healthy Adults, mTBI: mild Traumatic Brain injury patients, R1, R2, and R3 indicate the three predetermined dimensions used to plot participant cases through MDS.

For comparison, we computed a dissimilarity matrix of laplacian kernels using FCGs which were subsequently topologically filtered through conventional MST (Tewarie et al., 2015). Results of multidimensional scaling displayed in Fig. 4 reveal poor separation of individual cases especially among the healthy adult, mTBI, and typically achieving children. Classification accuracy did not exceed 45% for either the pairwise or the multi-group contrast.

**Fig. 4.**
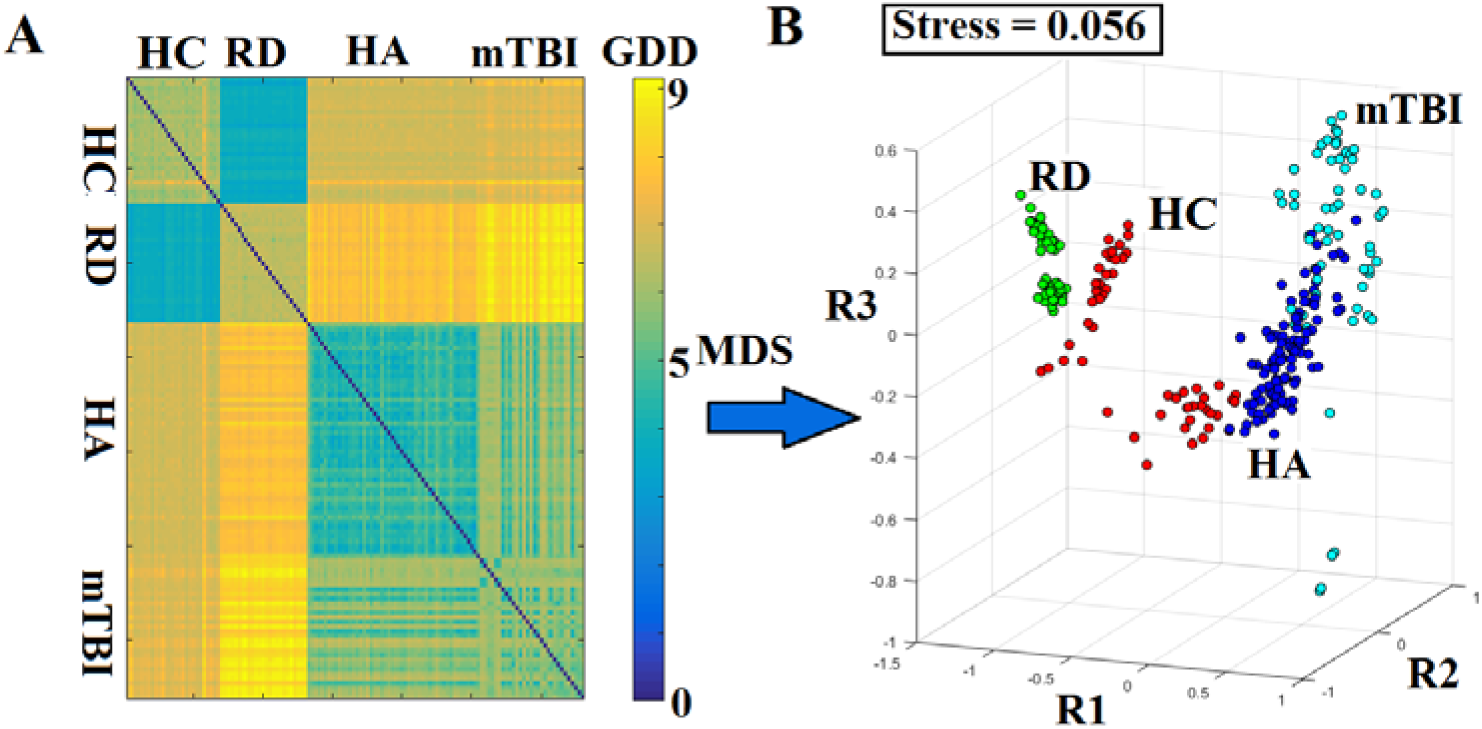
Topological filtering of Graph Diffusion Distance values using conventional MST. Dissimilarity matrix and Multi-dimensional Scaling (MDS) results for subject-specific FCGs (N=132) using the conventional method. All conventions are similar to those in Fig. 3.

To further ensure that discrimination between healthy adults and adults with a history of mTBI was not associated with differences in gender ratio between the two groups, the entire analysis was replicated on gender-matched subsets of the two groups (n=44 and n=28, respectively). Results presented in Section 4 of the Supplementary Material confirmed the superiority of topological filtering using OMST as compared to conventional MST in deriving FCGs that clearly distinguish between the two clinical groups.

### Group differences on Relative Power

Statistical filtering of relative power features derived a total of 14 features that were used for contrasting non-impaired and reading disabled readers resulting in an average of 67.3% correct classification (see Table S1 in Supplementary Material). Discrimination accuracy between non-impaired readers (children) and healthy young adults averaged 62.3% using a total of 49 features. Similarly, discrimination accuracy between mild traumatic brain injury (mTBI) subjects and healthy young adults averaged 68.1% using a total of 11 features. Finally, the multi-group discrimination accuracy did not exceed 53.3% using a total of 48 features.

### Group-specific Dominant Coupling Modes

Characteristic Dominant Coupling Modes, based on SMI, for each group of participants are displayed in the comodulograms of Fig. 5. Each 2D matrix tabulates the probability distribution of functional connections associated with intra-(diagonal cells) or inter-frequency coupling (cells above the diagonal). A notable finding is the higher proportion of significant cross-frequency coupling modes among typically-achieving students (12%; compared to only 5% in the reading-disabled group). Conversely, the latter group showed slightly higher within-frequency DICMs in the 0, a_1_, and β_1_ bands compared to non-impaired readers. Interestingly, both groups showed prominent DICMs in the δ band (Fig. 5A vs. Fig. 5B).

**Fig. 5.**
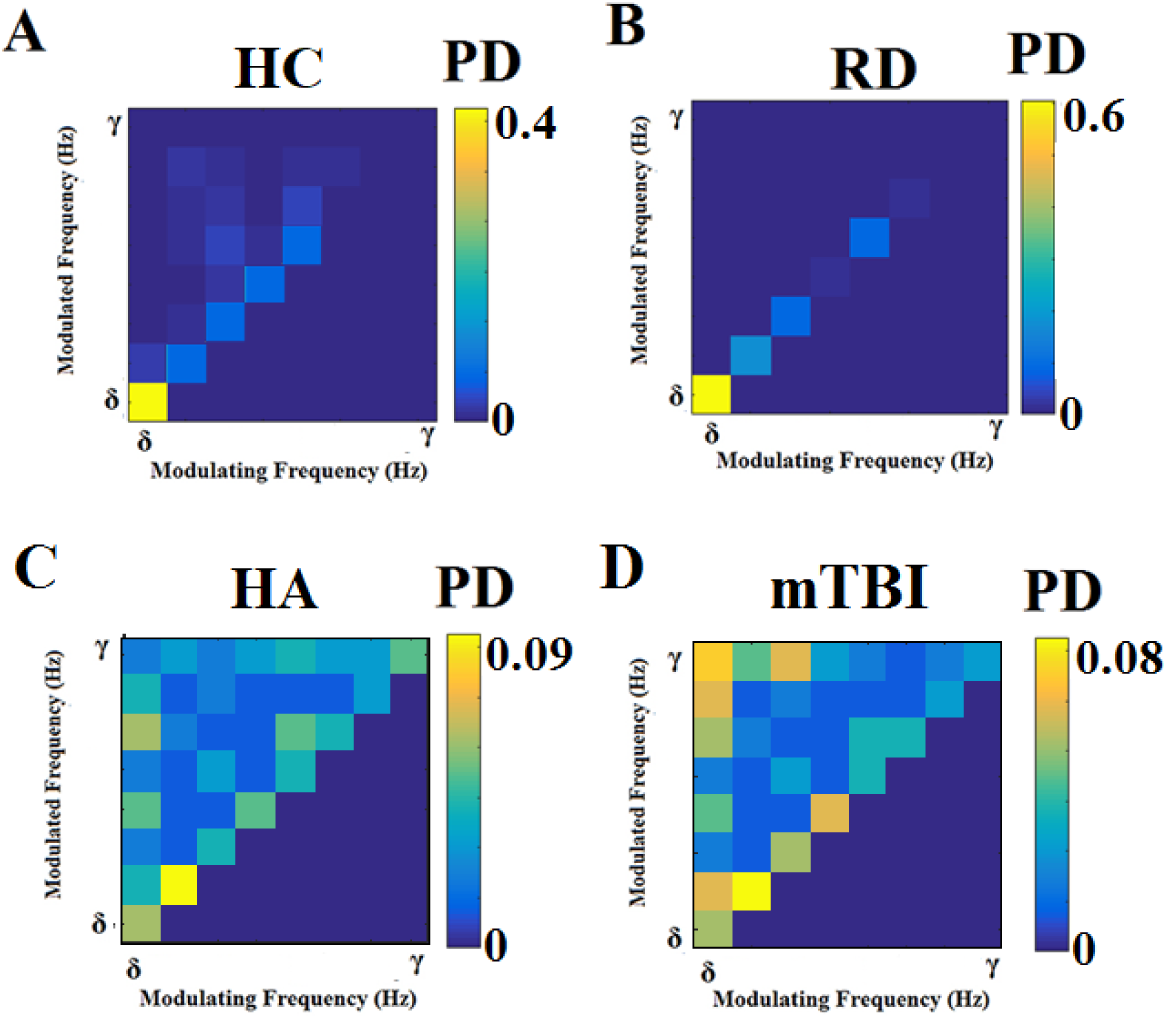
Group-averaged empirical Probability Distributions of dominant intrinsic coupling modes. Within-frequency coupling is shown in diagonal cells whereas cross-frequency interactions are shown in the off-diagonal cells. Abbreviations; HC: Typically developing children, RD: Reading-Disabled children, HA: Healthy Adults, mTBI: mild Traumatic Brain injury patients.

Moreover, the comodulograms of both adult groups were characterized by substantially higher relative contributions of cross-frequency DICMs as compared to groups of younger participants (with over 40% of DICMs representing cross-frequency interactions). Compared to healthy adults, participants with a history of mTBI displayed (i) prominent modulation of all higher frequency oscillations by δ frequencies and, (ii) more prominent within-frequency DICMs in the α_1_ and α_2_ bands.

## Discussion

### Topological filtering using OMST

The Orthogonal Minimal Spanning Tree (OMST) algorithm was introduced in the present work as a convenient, data-driven and computationally efficient method to perform topological filtering of a dense functional brain network. This method was introduced previously by our group in order to help identify the essensial connections within a given network in a manner that optimizes information flow between every element (node) of the network while mainitaining the functional/metabolic cost of connections at a minimum. Results from classification analyses on sensor-level MEG data from 132 participants confirmed the prediction that network metrics obtained from OMST-filtered networks would be sensitive to age-and diagnostic-group categories. Moreover, the superiority of OMST over the conventional MST approach, which is often too sparse to capture the most significant connections of a network, was unquestionable (100% vs. 45% classification accuracy in the multi-group analyses). The high classification accuracy based on the topologically filtered sensor networks via OMST underlines the effectiveness of this method to capture the most essential pathways of information flow within a given functional brain network. Robust group discrimination results were demonstrated for functional connectivity graphs derived from two complementary types of interdepenceny metrics: within-and cross-frequency coupling (Dimitriadis et al., 2015c; 2016a,b,c,d), as well as the phase lag index, which is considered to be less susceptible to volume conduction effects (Stam et al., 2007).

In the current application example, static sensor-level networks were analyzed, although the OMST method may be suitable for a variety of features (including dynamic metrics of MEG resting state data, and also network metrics derived from task-related MEG recordings; Dimitriadis et al., 2017). The OMST method was supplemented by estimation of diffusion distance metric (GDD), bearing distinct advantages over traditional approaches used to derive person-specific functional connectivity graphs, which rely on either unstructured data or vectors. In contrast, the diffusion distance metric was designed to identify systematic individual differences in functional brain networks, associated with distinct patterns of modeled information flow (Dimitriadis et al., 2015b). The GDD metric substituted Euclidean distance in a k-NN classifier as a more appropriate distance metric that respects the 2D format of the functional connectivity graph.

### Developmental and clinical correlates of OMST-derived functional networks

Although the present findings were based on sensor-level data, limiting their anatomic interpretability, the fact that we used planar-gradiometer neuromagnetic data to estimate patterns of within-and cross-frequency coupling modes permits certain conclusions regarding certain apparent features of underlying brain networks at rest. At a global network level, participants with a history of developmental (RD) or acute brain damage (mTBI) demonstrated lower global efficiency and higher diameter indices compared to typically achieving students and healthy adults, respectively (Fig. 2; Antonakakis et al., 2016; Dimitriadis et al., 2013, 2015a, 2016b). This finding is consistent with less efficient information flow during rest (at least at the sensor level) in both clinical groups compared to their age-matched typical/healthy counterparts. Compared to both student groups, adult comodulograms were also characterized by substantially higher proportions of cross-frequency interactions (40% and 45%, for young adults and mTBI patients, respectively). This finding is in accordance with the proposed trend toward more complex communication modes between remotely located neuronal assemblies with development (Buzsaki et al., 2012; Basar et al., 2016; Deco et al., 2017; Stamoulis et al., 2015). Interestingly, students who experienced severe reading difficulties in the present study were even less likely to display cross-frequency coupling modes at rest (5% of the total DICMs as compared to 12% among age-matched typical readers).

Moreover, history of mild traumatic brain injury was associated with a higher proportion of cross-frequency interactions involving δ modulating oscillators, in agreement with a recent report of abnormalities in functional brain networks in δ frequencies in mTBI (Dunkley et al., 2015).

### Limitations and future directions

The present study has several noteworthy limitations. Firstly, static networks were analyzed, although the OMST method is suitable for a variety of features (including dynamic metrics of MEG resting state data, and also network metrics derived from task-related MEG recordings). Secontly, connectivity patterns reflecting cross-frequency coupling were explored at the sensor-level. At this temporal scale, source localization (and related arbitrary choices of algorithms and anatomic templates) may introduce significant distortions to the source-level (reconstructed) signals. This added layer of complexity, although in principle desirable for enhancing the anatomic relevance of results, would very likely have confounded the primary goal of the study, namely to assess the capacity of OMST as a data-driven technique to derive sufficiently sparse graphs which could, in turn, reliably differentiate between age-and clinical diagnosis groups of participants. Thirdly, as presently applied OMST did not take into account the actual anatomic distance between sensors. Especially when applied to source-level data, this feature may enhance the sensitivity of the technique to explore functional cortical networks and can be aided by DTI-tractography data. Finally, independent assessment of the performance of the proposed algorithms and analysis pipeline on a new sample that includes both healthy participants and clinical groups is paramount in order to establish their generalizability.

## Conclusions

Orthogonal Spanning Trees is a promising method to identify important features (connections) of densely interconencted functional networks as represented by both conventional (PLI) and novel connectivity metrics (Symbolic Mutual Information). Integrating OMST-based network analyses with the notion of dominant coupling modes can offer complementary information regarding functional changes in the resting connectivity during the course of human development and also in relation to both developmental disorders and acute brain insults.

## Acknowledgements

This research was supported in part by grant P50 HD052117 from the Eunice Kennedy Shriver National Institute of Child Health and Human Development (NICHD). The content is solely the responsibility of the authors and does not necessarily represent the official views of the NICHD or the National Institutes of Health. SID was supported in part by MRC grant MR/K004360/1 (Behavioural and Neurophysiological Effects of Schizophrenia Risk Genes: A Multi-locus, Pathway Based Approach) and by a MARIE-CURIE COFUND EU-UK FELLOWSHIP.

## Notes

The Matlab code to implement OMST and the analysis pipeline described in this work is accessible at researchgate (https://www.researchgate.net/profile/Stavros_Dimitriadis), personal website (http://users.auth.gr/~stdimitr/software.html) and github (https://github.com/stdimitr/multi-group-analysis-OMST-GDD).

## Author Disclosure Statement

No competing financial interests exist.

